# Cell-free synthesis and purification of recombinant nucleocapsid (N), membrane (M), and envelope (E) proteins

**DOI:** 10.1101/2024.05.24.595851

**Authors:** Lin Wang, Mingming Fei, Wenhui Zhang, Sen Lin, Zhihui Jiang, Shun Zhang, Yao Wang

## Abstract

Rapid production of soluble recombinant antigens is important for developing *in vitro* diagnostics, vaccines, and drugs against virus such as severe acute respiratory syndrome coronavirus 2 (SARS-CoV-2). In this research, hard-to-express nucleocapsid, membrane, and envelope proteins were successfully expressed by an Escherichia coli-based cell-free protein synthesis system. The solubility of the proteins was optimized using various amphipathic molecules. Most of the impurities were easily removed by a one-step Ni-NTA affinity chromatography. This study provides an easy and quick alternative for virus’s trans-membrane and nucleotides associated recombinant protein expression, which has potential downstream application for early screening of newly emerging viruses.

## 1 Introduction

Coronavirus Disease 2019 (COVID-19) is an acute respiratory infection caused by severe acute respiratory syndrome coronavirus 2 (SARS-CoV-2) ^[1,2]^. Coronavirus contains positive-sense, single-stranded RNA ^[3]^. The virus can invade the respiratory, gastrointestinal, and central nervous system of humans, and birds, bats, mice, etc. Since the outbreak of COVID-19, the number of confirmed cases and deaths has continued to increase ^[4]^. Development of sensitive, rapid, and specific diagnostic testing technologies is needed for such pandemic at early stage.

There are at least 27 proteins in SARS-CoV-2. They include 15 non-structural proteins (nsp1∼nsp10, nsp12∼nsp16), four structural proteins (nucleocapsid, spike, membrane, and envelope proteins) and eight accessory proteins (3a, 3b, p6, 7a, 7b, 8b, 9b, and orf14) ^[5, 6]^. The 5’-end of the virus genome mainly encodes some of the non-structural proteins and the 3’-end encodes the structural proteins. As one of the main structural proteins of coronavirus, the spike (S) protein is key in virus particle adsorption and entry into host cells ^[7]^. The S protein amino acid sequence determines the specificity of coronavirus infection and the host range of the virus. The membrane (M) protein is the most abundant protein in the virus envelope. This determines the shape of the virus envelope ^[8, 9]^. It is the core of the coronavirus packaging and the main driving force for the formation of the coronavirus envelope ^[10]^. M protein interacts with all other structural proteins but cannot promote the formation of virus particles alone. The envelope (E) protein is the smallest structural protein ^[11]^. In the process of virus replication, the E protein is expressed in large quantities in infected cells, but only a small part is assembled into the virus envelope, which plays an important role in the process of virus particle generation and maturation^[12]^. Both M and E protein contain trans-membrane domains. The nucleocapsid (N) protein is located inside SARS-CoV-2. It is also an antagonist of interferon and a virus-encoded RNA interference inhibitor related to virus replication ^[13, 14]^.

The cell-free protein synthesis (CFPS) system enables rapid and efficient protein synthesis *in vitro* ^[15]^. The system mainly relies on mRNA or DNA (linear DNA or circular plasmids) as templates, and completes protein synthesis under the action of enzymes in cell extracts ^[16]^. Since the first use of CFPS to discover codons in 1961, CFPS applications have rapidly expanded and becoming an efficient research tool ^[17]^. Compared with the traditional cell-based expression system, the use of CFPS can overcome the physiological limitations of cells owing to its controllable operation and high tolerance to toxic proteins. This permits the *in vitro* expression of many complex proteins that are difficult to synthesize in the cells ^[18]^. CFPS-mediated expression of transmembrane proteins has been further facilitated by detergents and nanodiscs. Nanodiscs are discoidal bilayer membrane structures composed of membrane scaffold proteins (MSPs) and phospholipid molecules. Accumulating data have implicated nanodics as a potent cofactor to produce active transmembrane proteins ^[19]^. Pre-assembled nanodics with different lipid types and MSPs can be introduced into a standard CFPS reaction by direct mixing, making it a powerful tool for screening transmembrane protein expression conditions.

M, E, and N proteins are key antigens for coronavirus detection. In this study, we used an inexpensive CFPS system based on *E*.*coli* to express soluble M, E, and N proteins of SARS-CoV-2 to provide biological materials for *in vitro* diagnostics of COVID-19.

## 2. Materials and Methods

### 2.1 Synthesis of Target Sequences and Construction of Recombinant Vectors

Full-length DNA sequences of M, E, N, and S proteins of SARS-CoV-2 were obtained from NCBI (GenBank: MT745601.1) and codon optimized according to the codon preference of *E. coli*. The gene fragments were inserted into pET28a vector between the *Bam*HI and *Xho*I restriction sites.

### 2.2 CFPS

Standard CFPS reactions were performed using *E. coli* extracts, amino acid mix, and reaction mix as described by Tang *et al*. (2023)^[20]^. The reactions were performed at 30°C for 15 h. MSP1E3D1 1,2-dioleoyl-sn-glycero-3-phosphoglycerol (DOPG) (and MSP1E3D1 1-palmitoyl-2-oleoyl-sn-glycero-3-phosphoglycerol (POPG) nanodiscs (Zhengzhou Heang Biotech Co., Ltd) and detergent were added to 200 µL cell-free protein expression reactions as described in the text.

### 2.3 Protein Purification

CFPS sample supernatant was applied to Ni-NTA resin equilibrated with a binding buffer containing PBS (pH 7.5) at 4°C. The column was washed with a 10-column volume of buffer containing 50 mM imidazole in PBS (pH 7.5). The recombinant polyhistidine-tagged proteins were eluted with elution buffer containing 500 mM imidazole in PBS (pH 7.5). Finally, protein samples were concentrated by ultrafiltration using Amicon ultra centrifugal filter devices with a molecular weight cut-off of 10K (Millipore) and dialyzed in PBS (pH 7.5) using a disposable dialysis device (Zhengzhou Heang Biotech Co., Ltd). The purified proteins were analyzed by SDS-PAGE.

### 2.4 Identification of Recombinant Proteins

Protein samples were separated by 15% SDS-PAGE for 80 min at 100 V. For western blotting, the resolved proteins were transferred to polyvinylidene fluoride (PVDF) micro-porous membranes using the semi-dry iblotter (Invitrogen). Each membrane was blocked in 10 mL PBS containing Tween (PBST) and 5% skimmed milk powder for 2 h at room temperature (RT). The proteins were stained with an anti-His-6-tag rabbit antiserum (Cat. K200060M, Solarbio Co. Ltd.) diluted 1:1000 PBST 1:1000) and incubated at RT for 2 h. Subsequently, the membrane was washed three times with PBST for 5 min each time. Horseradish peroxidase (HRP)-labeled goat anti-rabbit IgG (Cat. AS003, ABclonal Co. Ltd.) diluted 1:2,000 was added as the secondary antibody. After incubation at RT for 2 h, the membrane was washed with PBST three times for 5 mins each time. Finally a color developing solution was applied ^[21]^.

## 3. Results

### 3.1 Expression and Solubiliy Analysis of Recombinant Proteins

The molecular weights of full-length SARS-CoV-2 M, E, N, and S proteins were 25, 9, 47, and 142 kD, respectively. M, E, and N proteins were successfully overexpressed using the CFPS system, while S protein was not (**Fig. 1**). The samples were centrifuged at 15000×g for 10 min and the supernatants were analyzed by SDS-PAGE. As the expressed proteins were not or only partially soluble detergent and nanotiscs were added in the same CFPS reactions. Solubilities of M and E proteins were significantly improved after adding detergent (**Fig. 2**). However, no improvement for N protein was observed when adding detergent or nanodiscs. Different conditions were applied for CFPS reactions of S protein, however expression was not improved.

**Fig. 1.**
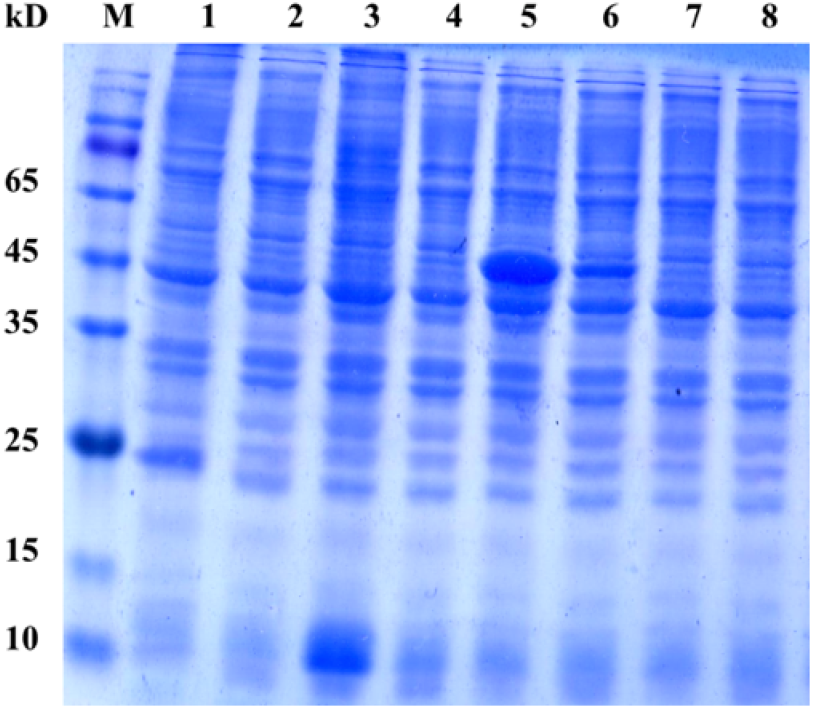
SDS-PAGE analysis of M, E, N, and S proteins expressed by standard CFPS reactions. Lane 1-8 are cell-free reaction samples of M protein (crude), M protein (supernatant), E protein (crude), E protein (supernatant), N protein (crude), N protein (supernatant), S protein (crude) and S protein (supernatant), respectively. Lane M contains molecular weight markers (Genstar Co. Ltd.).

**Fig. 2.**
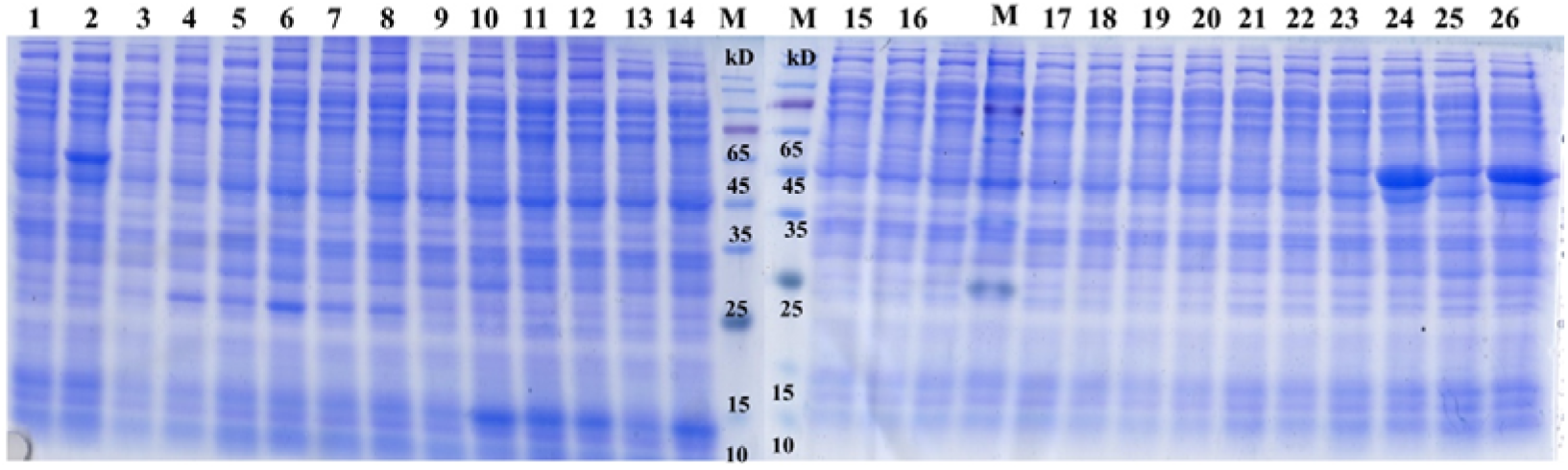
SDS-PAGE analysis of M, E, N, and S protein expressed by CFPS reactions with detergent or nanodiscs. Crude and supernatant sample analyzed in each lane are labeled as below. 1, supernatant N protein with detergent; 2, crude N protein with detergent; 3, supernatant M protein with MSP1E3D1 POPG; 4, crude M protein with MSP1E3D1 POPG; 5, supernatant M protein with MSP1E3D1 DOPG; 6, crude M protein with MSP1E3D1 DOPG; 7, supernatant M protein with detergent; 8, crude M protein with detergent; 9, supernatant E protein with MSP1E3D1 DOPG; 10, crude E protein with MSP1E3D1 DOPG; 11, supernatant E protein with detergent; 12, crude E protein with detergent; 13, supernatant E protein with MSP1E3D1 POPG; 14, crude E protein with MSP1E3D1 POPG; 15, supernatant S protein with detergent in 16°C; 16, crude S protein with detergent in 16°C; 17, supernatant S protein with detergent in 25°C; 18, crude S protein with detergent in 25°C; 19, supernatant S protein with detergent in 30°C; 20, crude S protein with detergent in 30°C; 21, supernatant S protein with detergent in 35°C; 22, crude S protein with detergent in 35°C; 23, supernatant N protein with MSP1E3D1 POPG; 24, crude N protein with MSP1E3D1 POPG; 25, supernatant N protein with MSP1E3D1 DOPG; and 26, crude N protein with MSP1E3D1 DOPG.

### 3.2 Purification of Recombinant Proteins

To produce larger quantities of recombinant proteins, M, E, and N proteins protein Ni-NTA resin was used. Results were analyzed by SDS-PAGE (**Fig. 3**).

**Fig. 3.**
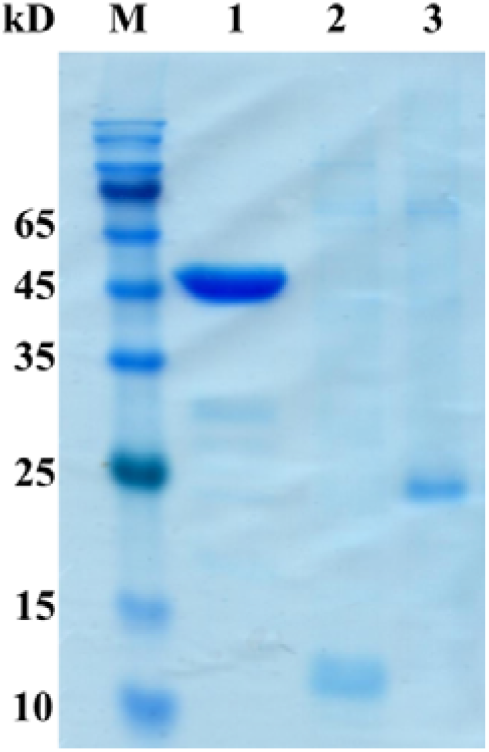
SDS-PAGE analysis for purified proteins. Lanes 1-3 contain N, E, and M protein, respectively. Lane M contains molecular weight markers (Genstar Co. Ltd.).

### 3.3 Recombinant Protein Identification

Western blotting using anti-His6 antibodies (**Fig 4**) demonstrated that N, M, and and E protein were successfully expressed and purified. The activities of the purified proteins were identified by ELISA and kinetic assays against their antibodies or binding partners and reported elsewhere.

**Fig. 4.**
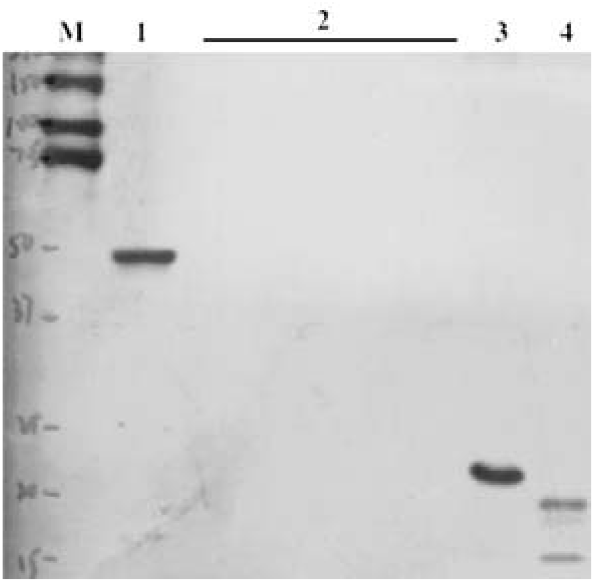
Western blotting of N, E, and M proteins. Lane 1, 3 and 4 are N, E and M proteins. The loading volume is 5, 0.5, and 0.5 _μ_g respectively. Lane 2 contains negative controls. Lane M contains molecular weight markers (Genstar Co. Ltd.).

## 4. Discussion

SARS-CoV-2 and Middle East respiratory syndrome coronavirus identified in 2012 belong to Beta-coronavirus genus ^[5]^. Their genomes share 80% sequence similarity with SARS-CoV, which is a typical coronavirus with enveloped single strand forward RNA ^[22]^. SARS-CoV-2 is mainly transmitted through droplets and close contact. After the virus enters the human body, it mainly replicates in the alveoli, causing the aggravation of inflammatory reaction in the lung and damaging the lung tissue. The main clinical manifestations are fever, fatigue, and dry cough. Some cases cause systemic inflammatory reaction which can be life-threatening ^[23]^.

The main structural components of coronavirus are M, E, and N proteins. N protein forms a viral ribonucleoprotein complex with 30 kb long viral genome RNA. E protein oligomerizes to form a “viral protein” ion channel. Even though M coordinates the assembly of virions, the interaction between M and E seems to be necessary for the formation of virions ^[24]^. After SARS-CoV-2 infection, specific IgM is detected as early as the third day of infection, and the peak appears 2 to 3 weeks after infection. Specific IgG antibodies can appear as early as 4 days and reaches a peak after 17 days ^[25, 26]^. An effective method for detecting SARS-CoV-2 IgM antibodies is a useful tool for identification of species of COVID-19, and for early diagnosis and treatment.

In this study, synthetic genes encoding SARS-CoV-2 M, N, E proteins were used to express the corresponding proteins through a detergent or nanodisc assisted CFPS system. The solubility of these hard-to-express proteins were significantly improved by adding detergent and nanodiscs in cell-free reactions. A single-step Ni^2+^ affinity chromatography was utilized to remove most of the impurities in the overexpressed protein samples after a short centrifugation. Unfortunately, S protein was not expressed despite using various physical and chemical conditions. The size of S protein (largest among the four structural proteins) and its complexity may have hindered overexpression. The use of eukaryotic cell-free systems may promote the overexpression of the S protein as cell organelles, chaperones, and codon preference may facilitate its overexpression. This study provides an easy and quick alternative for SARS-CoV-2 related recombinant protein expression, which has potential downstream application for early screening of viruses.

## Data Availability Statement

All data used to support the findings presented here are available upon request to the corresponding author.

## References

[1] E. Stokes, L. Zambrano, K. Anderson, E. Marder, K. Raz, S. Felix, Y. Tie, K. Fullerton, Coronavirus Disease 2019 Case Surveillance — United States, January 22–May 30, 2020. MMWR Morbidity and Mortality Weekly Report 69 (2020).

[2] H. Swann, A. Sharma, B. Preece, A. Peterson, C. Eldridge, D. Belnap, M. Vershinin, S. Saffarian, Minimal system for assembly of SARS-CoV-2 virus like particles. Scientific Reports 10 (2020).

[3] M. Batra, R. Tian, C. Zhang, E. Clarence, C. Sacher, J. Miranda, J. Fuente, M. Mathew, D. Green, S. Patel, M. Bastidas, S. Haddadi, M. Murthi, M. Gonzalez, S. Kambali, K. Santos, H. Asif, F. Modarresi, M. Faghihi, M. Mirsaeidi, Role of IgG against N-protein of SARS-CoV2 in COVID19 clinical outcomes. Scientific Reports 11 (2021).

[4] Z. Long, C. Wei, X. Dong, X. Li, H. Yang, H. Deng, X. Ma, S. Yin, Y. Qi, T. Bo, Simultaneous quantification of spike and nucleocapsid protein in inactivated COVID-19 vaccine bulk by liquid chromatography-tandem mass spectrometry. Journal of Chromatography B 1181 (2021) 122884.

[5] A. Wu, Y. Peng, B. Huang, X. Ding, X. Wang, P. Niu, J. Meng, Z. Zhaozhong, Z. Zhang, J. Wang, J. Sheng, L. Quan, Z. Xia, G. Cheng, T. Jiang, Genome Composition and Divergence of the Novel Coronavirus (2019-nCoV) Originating in China. Cell Host & Microbe 27 (2020).

[6] W. Khan, N. Khan, A. Mishra, S. Gupta, V. Bansode, D. Mehta, R. Bhambure, A. Rathore, Dimerization of SARS-CoV-2 nucleocapsid protein affects sensitivity of ELISA based diagnostics of COVID-19, 2021.

[7] R. Graham, R. Baric, Recombination, Reservoirs, and the Modular Spike: Mechanisms of Coronavirus Cross-Species Transmission. Journal of virology 84 (2009) 3134–3146.

[8] B.W. Neuman, G. Kiss, A.H. Kunding, D. Bhella, M.F. Baksh, S. Connelly, B. Droese, J.P. Klaus, S. Makino, S.G. Sawicki, S.G. Siddell, D.G. Stamou, I.A. Wilson, P. Kuhn, M.J. Buchmeier, A structural analysis of M protein in coronavirus assembly and morphology. Journal of Structural Biology 174 (2011) 11–22.

[9] K. Narayanan, A. Maeda, J. Maeda, S. Makino, Characterization of the Coronavirus M Protein and Nucleocapsid Interaction in Infected Cells. Journal of Virology 74 (2000).

[10] B.W. Neuman, G. Kiss, A.H. Kunding, D. Bhella, M.F. Baksh, S. Connelly, B. Droese, J.P. Klaus, S. Makino, S.G. Sawicki, S.G. Siddell, D.G. Stamou, I.A. Wilson, P. Kuhn, M.J. Buchmeier, A structural analysis of M protein in coronavirus assembly and morphology. Journal of Structural Biology (2011) 11–22.

[11] A. Mehregan, S. Perez-Conesa, Y. Zhuang, A. Elbahnsi, D. Pasini, E. Lindahl, R. Howard, C. Ulens, L. Delemotte, Biophysical characterization of the SARS-CoV-2 E protein, 2021.

[12] D. Schoeman, B. Fielding, Coronavirus envelope protein: Current knowledge. Virology Journal 16 (2019).

[13] V.M. Corman, O. Landt, M. Kaiser, R. Molenkamp, C. Drosten, Detection of 2019 novel coronavirus (2019-nCoV) by real-time RT-PCR. Eurosurveillance 25 (2020).

[14] V.A.J. Smits, E. Hernández-Carralero, M.C. Paz-Cabrera, E. Cabrera, Y. Hernández-Reyes, J.R. Hernández-Fernaud, D.A. Gillespie, E. Salido, M. Hernández-Porto, R. Freire, The Nucleocapsid protein triggers the main humoral immune response in COVID-19 patients. Biochemical and Biophysical Research Communications 543 (2021) 45–49.

[15] N. Colant, B. Melinek, J. Teneb, S. Goldrick, W. Rosenberg, S. Frank, D. Bracewell, A rational approach to improving titer in E. coli-based cell-free protein synthesis reactions. Biotechnology Progress 37 (2020).

[16] H. Nakano, T. Yamane, Cell-free protein synthesis systems. Biotechnology Advances 16 (1998) 367–384.

[17] M. Nirenberg, J. Matthaei, The Dependence of Cell Free Protein Synthesis in E. coli upon Naturally Occurring or Synthetic Polyribonucleotides. Proceedings of the National Academy of Sciences of the United States of America 47 (1961) 1588–1602.

[18] S.-M. Schinn, A. Broadbent, W.T. Bradley, B.C. Bundy, Protein synthesis directly from PCR: progress and applications of cell-free protein synthesis with linear DNA. New Biotechnology 33 (2016) 480–487.

[19] I. Denisov, S. Sligar, Nanodiscs for structural and functional studies of membrane proteins. Nature Structural & Molecular Biology 23 (2016) 481–486.

[20] Y. Tang, S. Ma, S. Lin, et al., Cell-free protein synthesis of CD1E and B2M protein and in vitro interaction, Protein Expr. Purif. 203 (2023) 106209.

[21] J. García-Cordero, J. Mendoza-Ramírez, D. Fernández-Benavides, D. Roa-Velazquez, J. Filisola-Villaseñor, S.P. Martínez-Frías, E.S. Sanchez-Salguero, C.E. Miguel-Rodríguez, J.L. Maravillas Montero, J.J. Torres-Ruiz, D. Gómez-Martín, L.S. Argumedo, E. Morales-Ríos, J.M. Alvarado-Orozco, L. Cedillo-Barrón, Recombinant Protein Expression and Purification of N, S1, and RBD of SARS-CoV-2 from Mammalian Cells and Their Potential Applications. 11 (2021) 1808.

[22] P. Zhou, X.-L. Yang, X.-G. Wang, B. Hu, L. Zhang, W. Zhang, H.-R. Si, Y. Zhu, B. Li, C.-L. Huang, H.-D. Chen, J. Chen, Y. Luo, H. Guo, R.-D. Jiang, M.-Q. Liu, Y. Chen, X.-R. Shen, X. Wang, X.-S. Zheng, K. Zhao, Q.-J. Chen, F. Deng, L.-L. Liu, B. Yan, F.-X. Zhan, Y.-Y. Wang, G.-F. Xiao, Z.-L. Shi, A pneumonia outbreak associated with a new coronavirus of probable bat origin. Nature 579 (2020) 270–273.

[23] D. Wang, B. Hu, C. Hu, F. Zhu, X. Liu, J. Zhang, B. Wang, H. Xiang, Z. Cheng, Y. Xiong, Y. Zhao, Y. Li, X. Wang, Z. Peng, Clinical Characteristics of 138 Hospitalized Patients With 2019 Novel Coronavirus–Infected Pneumonia in Wuhan, China. JAMA 323 (2020).

[24] B. Boson, V. Legros, B. Zhou, C. Mathieu, F.-L. Cosset, D. Lavillette, S. Denolly, The SARS-CoV-2 Envelope and Membrane proteins modulate maturation and retention of the Spike protein, allowing optimal formation of VLPs in presence of Nucleoprotein, 2020.

[25] Q.-X. Long, B.-Z. Liu, H.-J. Deng, G.-C. Wu, K. Deng, Y.-K. Chen, P. Liao, J.-F. Qiu, Y. Lin, X.-F. Cai, D.-Q. Wang, Y. Hu, J.-H. Ren, N. Tang, Y.-Y. Xu, L.-H. Yu, Z. Mo, F. Gong, X.-L. Zhang, W.-G. Tian, L. Hu, X.-X. Zhang, J.-L. Xiang, H.-X. Du, H.-W. Liu, C.-H. Lang, X.-H. Luo, S.-B. Wu, X.-P. Cui, Z. Zhou, M.-M. Zhu, J. Wang, C.-J. Xue, X.-F. Li, L. Wang, Z.-J. Li, K. Wang, C.-C. Niu, Q.-J. Yang, X.-J. Tang, Y. Zhang, X.-M. Liu, J.-J. Li, D.-C. Zhang, F. Zhang, P. Liu, J. Yuan, Q. Li, J.-L. Hu, J. Chen, A.-L. Huang, Antibody responses to SARS-CoV-2 in patients with COVID-19. Nature Medicine 26 (2020) 845–848.

[26] Q. Deng, G. Ye, Y. Pan, W. Xie, G. Yang, Z. Li, Y. Li, High Performance of SARS-Cov-2N Protein Antigen Chemiluminescence Immunoassay as Frontline Testing for Acute Phase COVID-19 Diagnosis: A Retrospective Cohort Study. Frontiers in Medicine 8 (2021).

